# Dual-color μ-LEDs integrated neural interface for multi-control optogenetic electrophysiology

**DOI:** 10.1101/2024.07.30.605927

**Authors:** Eunah Ko, Mihály Vöröslakos, György Buzsáki, Euisik Yoon

## Abstract

Over recent decades, optogenetics has become a pivotal technique for elucidating the functionality of neuronal circuits in living organisms. By genetically modifying specific cells within targeted tissues to respond to particular optical stimuli, researchers can achieve precise activation or inhibition of these cells. This capability enables detailed investigations of neural circuitry with unprecedented accuracy. However, there is a rising need for hardware that supports bidirectional control in conjunction with electrophysiological recording. A significant challenge in this domain is the compact integration of dual light sources and a recording system. This study addresses this challenge through the development of a novel microfabrication and assembly technique for embedding dual-color micro-LEDs and recording electrodes into a Michigan-type neural probe structure, designated as DuoLite (Dual-color micro-LEDs Integrated Neural-Interface Optoelectrode for Multi-Control Optogenetic Electrophysiology). We present two device variants: (a) a small-group and (b) a large-group cell-targeted design, each incorporating micro-LEDs with a minimal area of <100 μm^2^ for both red and blue light. The design and assembly techniques for integrating all three components within a shank width of <100 μm are thoroughly detailed, and the functionality of the devices is validated through in vivo experiments.

## Introduction

Optogenetics facilitates the manipulation of microbial opsins within neurons using light stimulation, thereby making these opsins light-sensitive and enabling the control of ion channels through specific wavelengths of light [1]. The technique’s advantages, including spatiotemporal precision and cell type specificity, are crucial for dissecting the functions of individual neurons within complex neuronal circuits [2, 3]. Moreover, optogenetics can be integrated with complementary technologies, such as electrophysiology, to provide deeper insights into neuronal circuitry by analyzing cell signals during perturbation events [4-6].

Recent advancements in optogenetics have aimed at enhancing its capabilities, notably through the diversification of opsins to cover a broader range of light wavelengths [3]. The availability of multiple opsin types permits the possibility of more sophisticated genetic manipulations beyond the traditional single opsin-single light spectrum approach [7-10]. Emerging research highlights the significance of dual-color optogenetics, which allows for experimental designs involving either a single cell group modified with two distinct opsins for bidirectional control [11, 17-19] or two separate cell groups each modified with different viral constructs [20]. This ability to achieve multi-control opens new possibilities for modulating and exploring neural circuitry with increased complexity and resolution.

To effectively manipulate opsins using different viral vectors, it is crucial that the light wavelength spectra targeting these vectors are sufficiently separated to minimize overlap. This separation is essential to avoid unintended stimulation of non-targeted opsins within overlapping wavelength ranges, which could result in unexpected experimental outcomes. For example, using a commercially available blue light LED with a peak wavelength of 463 nm alongside a green LED may be suboptimal due to their overlapping spectral range of approximately 475 to 500 nm [21]. Therefore, some studies have favored blue and red light sources, as their wavelengths are more distinct and less prone to overlap [11-13, 22].

Efforts have been made to develop hardware systems capable of supporting multidirectional experiments by incorporating dual light sources [12-13]. However, these systems often lack integrated recording capabilities for electrophysiological studies combined with optogenetics, complicating cellular analysis. One approach demonstrated the feasibility of integrating two distinct light sources above recording electrodes using waveguide technology [11]. Nonetheless, this method encounters challenges due to the large footprint of the light delivery system and difficulties in integrating multiple light sources on a single probe shank. Ideally, a system would feature a scalable light source at the neural interface along with an integrated neural activity recording system, enabling simultaneous multi-color applications.

This work introduces an innovative microfabrication process that addresses these needs. The process involves: 1) fabricating a polyimide-encapsulated red micro-LED flexible probe and 2) designing and integrating this flexible red probe onto commercially available silicon-based blue-LED optoelectrodes [7], or onto other customized blue-LED optoelectrodes for small and large area targeted designs. The red micro-LEDs have a minimum size of 14 μm × 17 μm, showcasing their scalability. The wavelength separation between the blue and red LEDs remains reliable after full assembly. The characteristics of this dual-color probe, termed DuoLitE, will be detailed, and its performance will be validated through in vivo experiments involving head-fixed mice targeting the hippocampal region.

## Materials and methods

### Novel process architecture of the flexible red micro-LED probe

The red micro-LED flexible probe is constructed from a GaAs-based LED epitaxy wafer sourced from Powerway GaAs Wafers (Xiamen Powerway Advanced Material Co.). This wafer features a 350 μm thick GaAs base with an epitaxy layer approximately 6 μm thick. The epitaxy layer consists of n-type materials such as Si-doped GaInP or GaAs for the n-contact layer, a multi-quantum well (MQW) layer situated between the p- and n-type materials, and GaP or AlInP for the p-contact and p-type semiconductor layers at the top.

The fabrication process for the flexible red micro-LED probe involves nine masking steps. Initially, the wafer is cleaned with acetone and isopropanol (IPA) for a total of 10 minutes, followed by a 1-minute cleaning with an HCl solution, thorough rinsing with deionized (DI) water, and drying. The p-contact metal deposition is then performed by sputtering indium tin oxide (ITO) material (Lab 18-01 sputter, LNF) to a thickness of 200 nm, followed by annealing the wafer in rapid thermal annealing (RTA) equipment (Jetfirst 150, LNF) at 400°C in an argon (Ar) atmosphere for 5 minutes.

The first mask process defines the alignment marks and delineates the p-GaP contact metal extension, which will later connect to the metal traces. S1813 photoresist (PR) is applied and soft-baked at 90°C for 2 minutes. The mask is then exposed to the wafer using a stepper (GCA AS200 AutoStep, LNF) for 0.24 seconds. After developing the wafer in AZ726 developer, evaporation is carried out (Enerjet, LNF) to deposit a total of 100 nm of titanium and gold. The photoresist is then lifted off using a solvent bench.

Following the first masking process, the next step is to define the red LED mesas. The epitaxy layer, approximately 6 μm thick, is etched using a physical method to avoid the formation of chemical traps on the mesa sidewalls, which can degrade the performance of red LEDs, particularly their light output power [23]. A soft mask is used in this process. SPR 220 3.0 photoresist is spun and soft-baked on the wafer at 1500 rpm and 115°C for 2 minutes. The wafer is then exposed through a stepper for 0.6 seconds. After a post-exposure bake (PEB) at 115°C, the wafer is developed with AZ726 developer to achieve a photoresist thickness of around 4 μm. The wafer is then prepared for mesa etching using an ion milling machine (Nanoquest II Ion Mill, LNF). The mesa is etched slowly over approximately 2 hours, with an average etch rate of 50 nm/min. Once mesa etching is complete, the remaining photoresist is removed using PRS2000 chemical (see Fig. 3A-B, and K).

**Figure 1.**
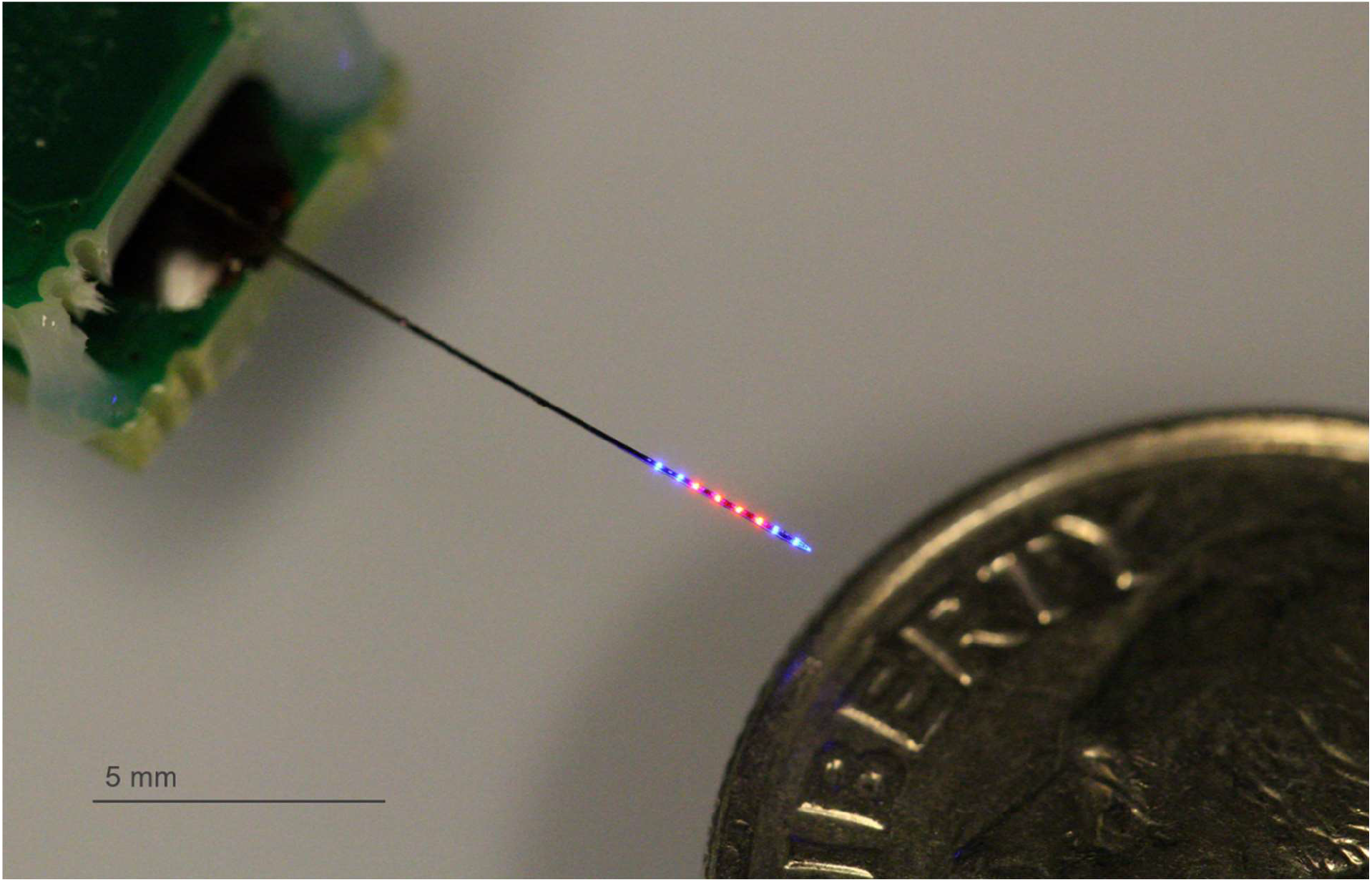
Bird’s eye view of DuoLite for deep-brain multi-control optogenetic electrophysiology. The DuoLite device is designed to conduct the roll of deep-brain neural interface with multi-controlled optogenetics through two distinct type of light source (463 nm *λ*_peak_ blue and 647 nm *λ*_peak_ red) with electrophysiology. Both LEDs and recording electrode sites are packed within the < 100 μm width of probe shank, where size and the spacing of LEDs and RECs are customizable depending on the purpose of the designed experiment. The specific type shown in this figure is defined as type B in this study, with 10 blue and red LEDs and 10 RECs are integrated in a 85 μm wide shank with 2 mm of top-down span.

**Figure 2.**
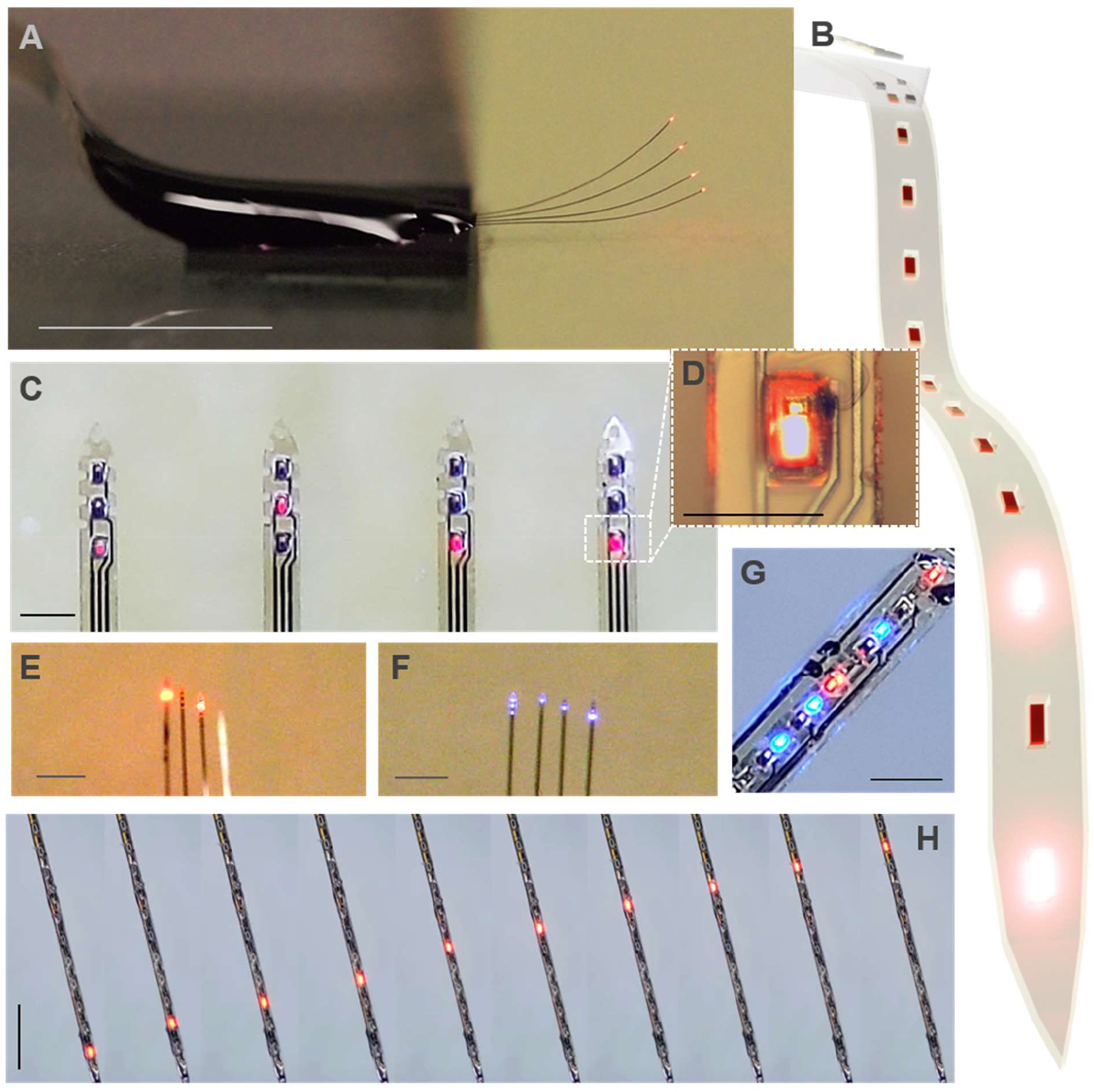
The red micro-LED integrated flexible neural interface (R-flex). **A)** Oblique view of R-flex, multi-shank type. Scale bar: 5 mm. **B)** Schematic of flexible red-μLED neural interface. **C)** Expanded view of **A)**, where the red-μLEDs are placed (on the tip side of the probe shanks). Scale bar: 100 μm. **D)** Expanded view of one red-μLED from **C)**. Scale bar: 50 μm. **E)** Top-view of R-flex in **A)**. Scale bar: 500 μm. **F)** Top-view of blue LED integrated flexible probe. Scale bar: 500 μm. **G)** Example stacked device composed of flexible red and blue-μLED probes. Scale bar: 100 μm. **H)** Individually functioning red-μLEDs in R-flex device. Scale bar: 500 μm.

**Figure 3.**
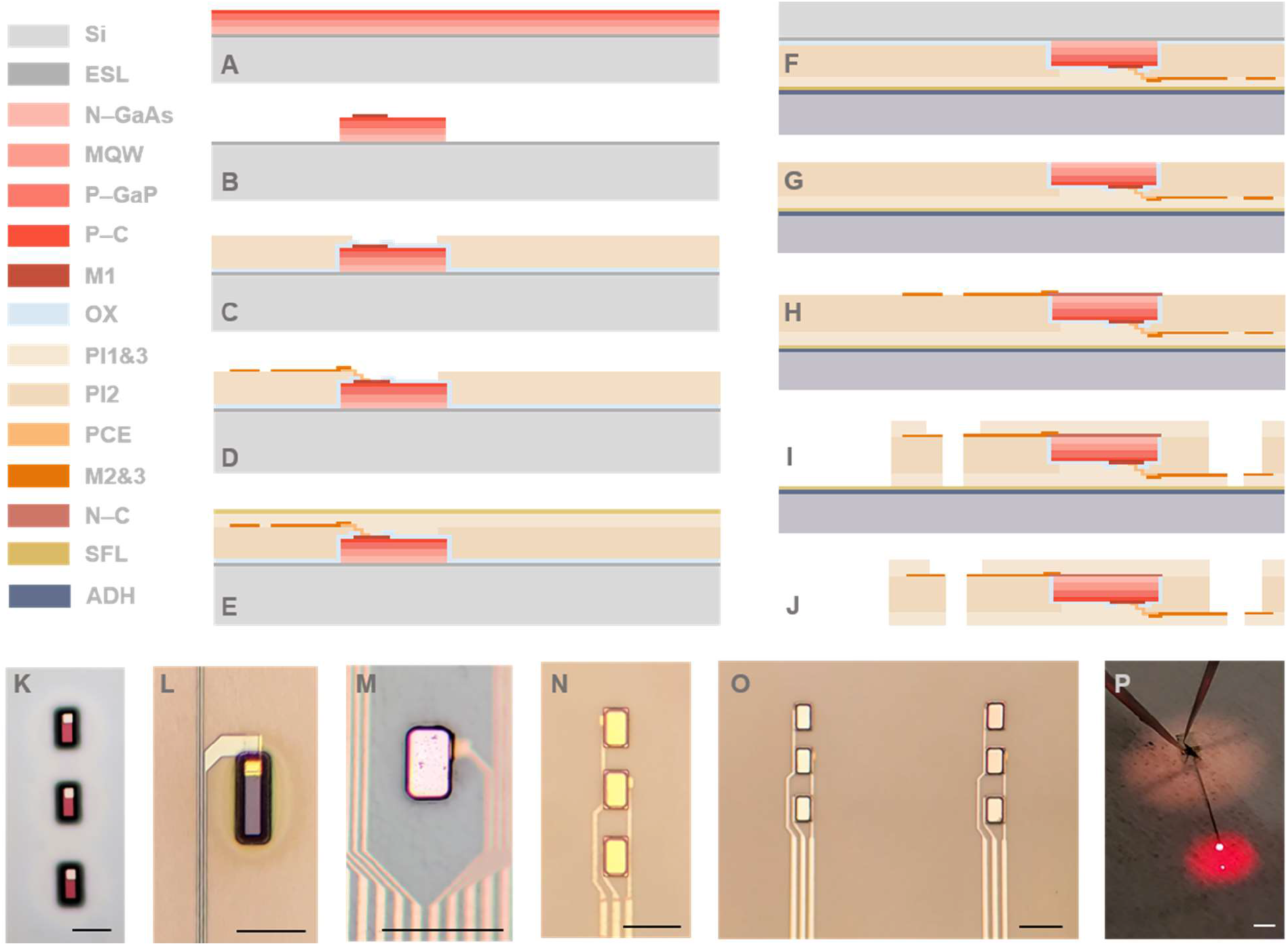
Fabrication process of red micro-LED integrated flexible neural interface (R-flex). A-J) side-view and K-P) top-view of the process. **A)** Starting point of the red-epitaxy wafer composed of silicon, n-and p-type layers and MQW layers. **B)** P-contact and M1 process followed by LED mesa definition. **C)** Mesa passivation, PI #1 and opening process. **D)** P-contact extension and metal line 1 process. **E)** PI #2 and sacrificial layer (SFL). **F)** Wafer transfer to the carrier wafer through adhesive pressure/temperature assisted bonding. **G)** Backside field removal (GaAs, etch stop layer (ESL), and oxide). **H)** N-contact and metal line 2 processes. **I)** PI #3 definition, probe outline etching. **J)** Release probe. The top view of the processes corresponds to the side view as follows; K) to B), L) to D), M) to G), N) and O) to H), P) to J). Scale bar 50 μm from K) to O), and 1 mm in P).

The LED mesas are tested after photoresist removal using a probe station, making contact with the mask 1 processed p-GaP contact extension and the bottom GaAs contact. After confirming the functionality of the LEDs, the wafer is passivated with an oxide layer approximately 550 nm thick, composed of Al2O3 and SiO2, deposited using Oxford ALD and P5000 PECVD (LNF). The first polyimide layer is then spun and cured to achieve a thickness of approximately 5.5 μm (PI2611, HD Microsystems, spin rate of 3000 rpm). Due to the significant step height difference between the LED mesas and the surrounding field region (6 μm), the polyimide layer is designed to match the mesa height, while the polyimide atop the mesas is subsequently etched away, and the wafer is planarized.

The next step involves etching away the polyimide on top of the mesas (mask 3). SPR 220 3.0 photoresist is spun and soft-baked at 1000 rpm and 115°C, then exposed using a stepper for 0.7 seconds. After a post-exposure bake (PEB), the wafer is developed for 50 seconds using AZ726. Subsequently, the polyimide on top of the mesas is etched using a Plasmatherm system (LNF) for approximately 17 minutes, achieving complete removal with an etch rate of 200 nm/min. Careful layout design is crucial to account for the slope of the photoresist pattern after development; a tight layout margin between the mesa edge and the photoresist edge may lead to unwanted etching of the polyimide on the mesa sidewalls. After polyimide etching, any remaining photoresist is thoroughly removed using acetone.

The next step, mask 4, involves opening the oxide layer on top of the P-GaP metal extension pattern (via) to facilitate electrical connection to the metal traces. Again, SPR 220 3.0 is spun at 1000 rpm and soft-baked, followed by a 0.7-second exposure. The wafer is then developed, and the oxide layers are etched away using reactive ion etching (RIE) equipment (LAM 9400, LNF). After treating the opened vias with an acid solution (HF) for thorough cleaning, the photoresist is stripped away.

The Mask 5 process addresses the step height difference between the top of the p-GaP metal and its extension (from mask 1) relative to the topmost layer of polyimide. Double-layer lithography is employed using LOR and SPR, with each layer spun at 1000 rpm. The photoresist layers are exposed for 0.7 seconds in the stepper and developed for 70 seconds. The wafer then undergoes treatment with oxygen plasma to remove residual organics, followed by sputtering of titanium and gold (totaling 300 nm, Lab 18-02, LNF). The double-layer photoresist is subsequently removed using Remover PG.

For the deposition of metal traces, the wafer is initially cleaned for 60 seconds in oxygen plasma to ensure the polyimide surface is clean and to enhance adhesion between the polyimide and metal. SPR is spun and baked at 4000 rpm and 115°C, then exposed for 0.4 seconds in the stepper. After a post-exposure bake, it is developed with AZ726 for approximately 40 seconds. Titanium and gold layers are then evaporated to a total thickness of 200 nm and lifted off using acetone. The wafer is now ready for testing the LEDs to assess the approximate yield of the p-metal traces (see Fig. 3D and L), marking the completion of this stage and preparation for subsequent processes.

Subsequent processes involve fully passivating the LEDs and traces with polyimide, which requires transferring the wafer and removing the substrate, as well as forming n-GaAs contact and traces for grounding. A second polyimide layer is coated onto the wafer to achieve a thickness of 2 μm. Next, a sacrificial layer is sputtered onto the wafer, which will later be removed to release the probe. Both the carrier wafer (sapphire) and the device wafer undergo spinning and curing of HD3007 at a spin rate of 3000 rpm and an oven temperature of 350°C. The two wafers are then bonded using a wafer bonder (EVG 510, LNF), applying a force of 1600 N at 350°C (see Fig. 3E and F).

The backside of the GaAs substrate is now exposed. Before etching the GaAs substrate, the surface is cleaned with oxygen plasma etching to remove any remaining organics. The wafer is then subjected to three steps of wet etching at the acid bench. First, the oxide is removed with BHF etching, followed by etching of the GaAs using an NH4OH:H2O2 solution for approximately 3 hours, exposing the etch-stop layer.

Subsequently, the etch-stop layer is removed using an H_3_PO_4_:HCl:O_2_ solution, which exposes the n-GaAs contact material (see Fig. 3G and M). Precise calculation of the etch rate and etching time is crucial during these wet etching steps, as even minor over-etching can damage the mesa sidewalls and remaining mesa islands, leading to degradation of the LED characteristics.

The wafer is then prepared for the Mask 7 process, which involves defining the n-GaAs contact metal. Double-layer lithography is employed using LOR and SPR at spin rates of 3000 rpm and 2000 rpm, respectively. The photoresist is exposed for 0.6 seconds, followed by a post-exposure bake (PEB), and then developed for 50 seconds. A stack of germanium, nickel, and gold is evaporated to a total thickness of 150 nm and then lifted off. The wafer surface is subsequently cleaned using oxygen plasma treatment.

The next step is defining the n-metal traces (Mask 8). SPR 220 photoresist is spun and baked at 4000 rpm and 115°C, exposed in the stepper for 0.4 seconds, and then subjected to a post-exposure bake (PEB). It is developed for 37.5 seconds using AZ726 developer, followed by 1 minute of oxygen plasma treatment. Titanium and gold layers, totaling 200 nm in thickness, are then evaporated. The metal trace pattern is formed through lift-off using acetone.

Finally, the backside of the wafer is covered with a 2 μm thick polyimide layer. Using an Al_2_O_3_ hard mask, the probe outline is etched for 40 minutes under reactive ion etching (RIE), and the probe is released by removing the sacrificial layer, making it ready for packaging (see Fig. 3I, J, and P).

### Fabrication of the compatible blue micro-LED optoelectrode (Large area blue optoelectrode)

Before proceeding to the DuoLite assembly, a brief explanation of the fabrication process for the large-area targeting blue silicon optoelectrodes, which are compatible with the large-area designed red LED probe (see Fig. 4E), is provided. The large-area optoelectrode is designed to incorporate blue micro-LEDs and recording electrodes (RECs) that are five times larger than those used in the MINT probe. The fabrication process architecture is similar to that of the MINT probe [7, 24], with the large-area targeting device featuring a 10 mm long shank, which includes a single shank design with 12 LEDs and 12 RECs, and a minimum trace pitch of 2 μm. Therefore, a high-density lithography process was employed for the probe metal traces [25]. Additionally, to minimize stress induced by the 1.5 μm thick oxide deposition upon process completion, SiO_2_ deposition was replaced with oxynitride deposition (GSI PECVD, LNF), reducing stress from approximately -200 MPa to around -59 MPa. This reduction in stress aims to minimize bending of the shank after release, which is crucial for reliable surgical implantation, as a bent shank would compromise precise probe placement. The target thickness of the shank was set at approximately 70 μm, controlled using deep reactive ion etching (DRIE) equipment (Pegasus 6, LNF).

**Figure 4.**
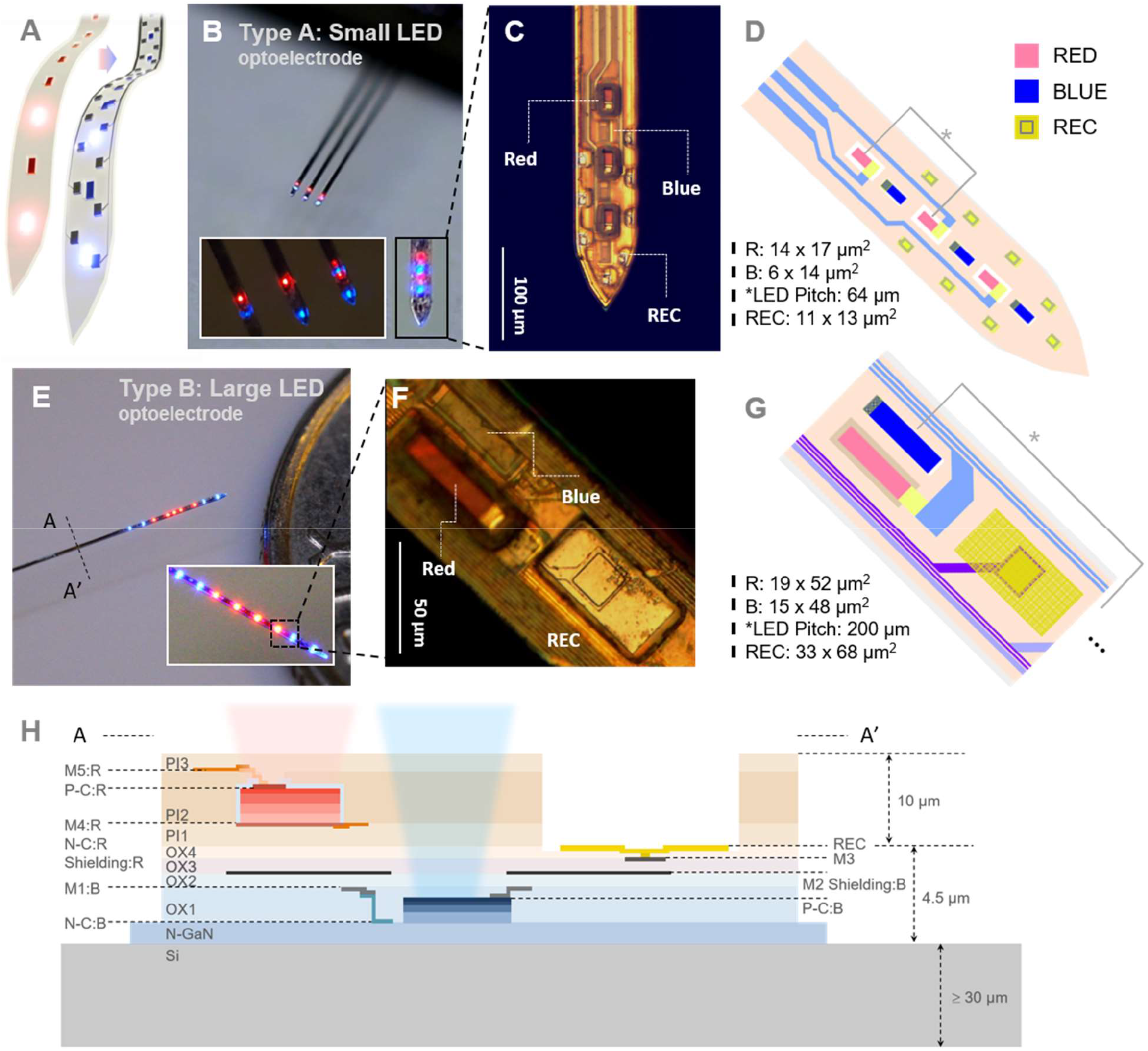
Design of DuoLite considering the placement of dual color micro-LEDs and recording electrodes. **A)** Schematic of stacking red micro-LED probe on top of blue micro-LED optoelectrode. **B)** Bird’s eye view of small-area targeting DuoLite. **C)** Expanded view of a single shank in figure B). **D)** Schematic image based on the designed layout of the stacked view of small-area targeting DuoLite shown in C). LEDs and recording sites dimensions are noted (R: area of red micro-LED and B: area of blue micro-LED) **E)** Bird’s eye view of large-area targeting DuoLite. **F)** Expanded view of the shank shown in inset figure of E). **G)** Schematic image based on the designed layout of the stacked view of large-area targeting DuoLite shown in F). **H)** Schematic of the cross-sectional view from A to A’ direction shown in figure E).

### Layout considerations and assembly

The design of the red micro-LED flexible probe is consistent with that of the blue micro-LED optoelectrodes used in both the MINT and large-area configurations. Recording electrodes are integrated into the silicon probe, with the red probe incorporating openings at its base to accommodate these recording electrodes, as illustrated in Figures 4B-G. In the small-area DuoLite configuration, micro-LEDs are arranged vertically, with each color interleaved between the other. Despite the top of the blue LEDs being covered with polyimide for the flexible red LED probe, no significant degradation in the characteristics of the blue LEDs was observed during in-vivo testing.

For the large-area targeting DuoLite, a custom-designed blue-LED optoelectrode serves as the base, with the flexible red probe stacked on top. In this configuration, the blue and red micro-LEDs are positioned side by side, with recording electrodes placed between the two types of LEDs. This arrangement is repeated throughout the probe design. Details regarding the LED area, LED pitch, and REC size are provided in Figure 4.

To accommodate potential misalignment, the polyimide openings for the recording electrodes are designed to be greater than 2 μm larger than the actual REC size. This design ensures that neither the blue nor red micro-LEDs, nor the REC sites, experience degradation, as will be discussed later. Figure 4H presents a cross-sectional schematic view along direction A to A’, as indicated in Figure 4E, offering a comprehensive view of all layers within the DuoLite structure. The overall device layout is depicted in Figure 1.

The entire structure is built upon a silicon base approximately 30 to 70 μm thick, with the blue LED optoelectrodes consisting of the LEDs, shielding layers, and recording electrodes. The top layer of the red LED flexible probe includes a bottom ground metal layer, which also functions as a shield, and a top Vin layer that drives the red micro-LEDs. Each type of LED is driven independently via connectors at the backend, where two stacked PCBs are secured with epoxy (see Fig. 1).

### In-vivo experiment

All animal procedures were approved by the New York University Animal care and Facilities committee. A female VGAT-Cre mouse (B6J.129S6(FVB)-Slc32a1tm2(cre)Lowl/MwarJ; JAX Labs, Maine) was injected in the CA1 region of the hippocampus (coordinate, in mm from bregma: −2.0 posterior, 1.5 mm right) with two AAVs, encoding Cre dependent ChrimsonR (AAV5-hSyn-FLEX-ChrimsonR-tdTomato) and CaMKII promoter driven ChR2 (AAV5-CaMKIIa-hChR2(H134R)-EYFP), resulting in expression of ChrimsonR in interneurons and ChR2 in pyramidal neurons (North Carolina Vector Core (UNC REF)). The mouse was additionally implanted with 3D-printed headpost [27] and stainless steel ground screw placed above the cerebellum. The mouse was habituated to head fixation over the course of one week. After habituation, the animal was head-fixed, and the electrode was lowered to dorsal CA1. Baseline recording was obtained, after which stimulation with 463.2 and 646.7 nm light was made at different intensities.

Current-controlled stimulation was used to drive individual μLEDs (OSC1Lite, 12-ch current source) [25]. Each color was controlled by an OSC1Lite. Neural data were acquired at 20 kHz using an Intan RHD2000 recording system. Offline, spikes were detected and automatically sorted using the Kilosort algorithm [28] followed by manual curation using Phy. Analysis was performed in MATLAB using custom scripts. To measure the effect of light stimulation on spiking activity, peristimulus time histograms (PSTHs) were built around stimulus onset (spike trains were binned into 10-ms bins). Baseline and light-induced spiking rate were calculated for each single unit. Baseline was defined as light-free epochs (2 s) between trials and stimulation period as the 463.2 and 646.7 nm light was on (200 ms). Wilcoxon-signed rank test was used to compare the mean firing rate per trial during baseline and LED stimulation.

## Result

### RED Micro-LED characteristics after wafer process

Figure 5 illustrates the measured characteristics of both large-area and small-area targeting DuoLite devices following full assembly and PCB fixation. As previously noted, it is crucial to ensure complete separation of the light illumination spectra for each type of micro-LED. This separation is essential because spectral overlap could lead to unintended cellular reactions. The injected viruses and genetically modified proteins in neuronal cells respond to a broader bandwidth rather than a very narrow spectrum [29]. The measured spectra show clear separation between the red and blue light emitted by each type of LED (see Fig. 5A).

**Figure 5.**
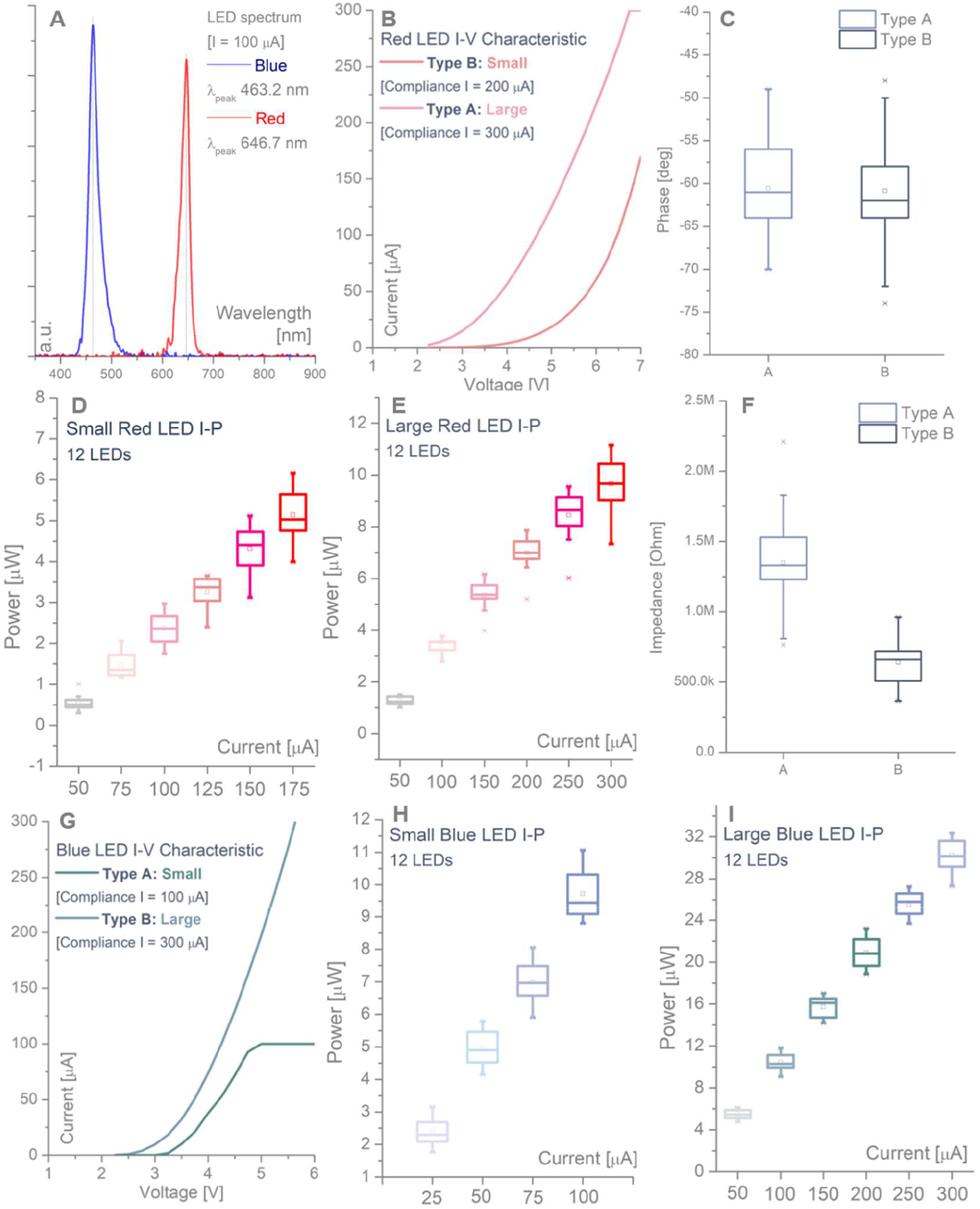
Device characteristics for type A and B DuoLite. **A)** Wavelength spectrum measured for blue and red micro-LEDs after full assembly of DuoLite. **B)** Current (I) – voltage (V) characteristics for small (type A) and large (type B) red micro-LEDs. **C)** Phase of the recording electrodes measured after full assembly of DuoLite for both types of devices. **D) and E)** Current (I) – optical power (μW) measured for type A and B DuoLite devices, respectively. **F)** Impedance magnitude (Ohm) value measured for both types of DuoLite devices after full assembly. **G)** Current (I) – voltage (V) characteristics for blue micro-LEDs on type A and B DuoLite. **H) and I)** Current (I) – optical power (μW) characteristics for type A and B blue micro-LEDs in DuoLite device.

Figure 5B depicts the current-voltage characteristics of the red micro-LEDs for both large and small configurations. The device is designed to achieve a power output greater than 1.5 μW within a voltage input range of up to 7 V, with the current characteristics being suitable for in-vivo applications. The light output power (μW) versus current (μA) is shown in Figures 5D and 5E for the small and large LEDs, respectively.

### Multicolor device characteristic after assembly

The red flexible probe was assembled and then aligned with the pre-processed blue LED optoelectrode. Following full assembly, the characteristics of the blue micro-LEDs were measured, as illustrated in Figures 5G, H, and I. Figure 5G shows the current versus voltage characteristics, revealing a slightly increased threshold voltage for the small-area targeting LEDs. This discrepancy is attributed to the microfabrication process, where larger LEDs feature larger oxide and polyimide opening contact via patterns, resulting in lower contact resistance after processing. Figures 5H and I display the optical power (μW) versus current (μA), demonstrating that there is no significant degradation in the absolute light power compared to previously reported blue-LED optoelectrodes [7]. This indicates that the additional layer of approximately 10 μm of polyimide does not substantially reduce the light power emitted by the LEDs.

Impedance characteristics are shown in Figures 5C and F. Figure 5C presents the phase value for the recording electrodes, while Figure 5F displays the impedance magnitude. Data were collected from a total of 50 sample microelectrodes, with Type A representing the small-area targeted device and Type B the large-area targeted device (REC area for Type A is 11×13 μm^2^, and for Type B is 33×68 μm^2^). The impedance characteristics are suitable for electrophysiological applications and will be used in in-vivo experiments. For these experiments, the Type A (small-area device) is employed for targeting the hippocampus area.

### In-vivo validation

The assembled dual color μLED probe (small-area targeting DuoLite), containing 32 recording sites and 12 blue and red μLEDs, was used to record neural activity from dorsal CA1 of an awake VGAT-Cre mouse (n = 1). The mouse was injected with a mixture of two AAVs, driving the expression of ChR2 under the CaMKII promoter (in pyramidal neurons, PYR) and ChrimsonR under the VGAT promoter (in interneurons, INT, Fig. 6A, B).

**Figure 6.**
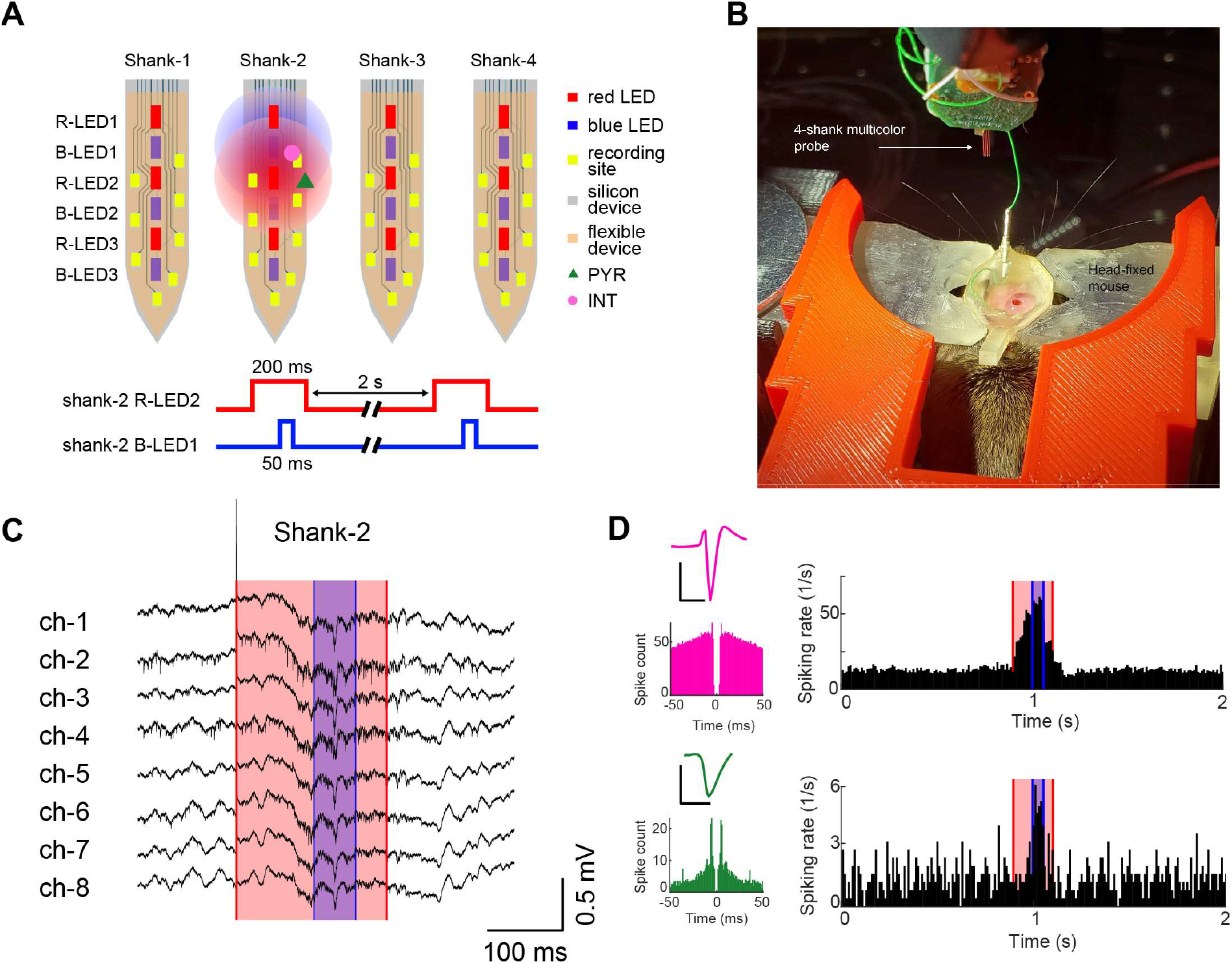
In-vivo validation of the DuoLite device type A. **A)** Schematic of four shanks with blue and red micro-LEDs and recording electrodes (top panel). Bottom panel shows the pulse pattern used for driving both color micro-LEDs. Both micro-LEDs are controlled by current where for red is 90 μA, and blue is 10 μA. **B)** Picture of the head-fixed experiment showing the 4-shank DuoLite device before implantation. **C)** Raw signal recorded from type A DuoLite device shank-2 used in head-fixed mouse experiment. **D)** Top panel shows the interneuron controlled through the red micro-LED (location shown as pink circle in figure B) where the R_LED 2 in shank 2 was used) and the bottom panel shows the pyramidal neuron controlled through the blue micro-LED (location shown as green triangle in figure B) where B-LED1 was used for stimulation).

Spontaneous neural activity was recorded on all shanks while light-induced neuronal activity was observed on the illuminated shank (shank-2 red-LED2 and blue-LED1, Fig. 6C). The stimulation artifact induced by the red LED was 1.88 ± 0.11 mV (mean ± SEM during LED onset) and 1.45 ± 0.09 mV (offset). As expected, the blue LED did not induce observable artifacts in the neural signal (11.6 ± 0.4 μV during onset and 9.9 ± 0.39 μV during offset; mean ± SEM). Illumination with 463.2 nm light pulses (50 ms, 10 μA, n = 234 trials) elicited spiking in 26/8 (30%) PYR and in 1/1 (100%) INT. 646.7 nm light pulses (200 ms, 90 μA, n = 234 trials) changed the spiking of a putative interneuron recorded on shank-2 (pink cell, Fig. 6 D). This putative single unit increased spiking during 635 nm light illumination (19.96 Hz baseline firing rate increased to 37.52 Hz; p < 0.05; Wilcoxon signed rank test), presumed to result from direct ChrimsonR-mediated depolarization. The same cell also increased its spiking during 463.2 nm illumination (mean firing rate is 52.71 Hz), presumed to result from synaptic inputs from the population of ChR2 expressing pyramidal neurons, which are known to make excitatory synapses on these neurons (Fig. 6 D). An example PYR responding to 405 nm light (mean firing rate is 1.3 Hz without and 6.41 Hz with stimulation, p < 0.05; Wilcoxon signed rank test) and having no response to 635 nm light is shown in Fig. 6 D (green cell).

## Discussion

The novel multicolor device is demonstrated by stacking a flexible red micro-LED probe onto an existing silicon-based blue-LED optoelectrode or a custom-designed silicon optoelectrode. The process begins with outlining and demonstrating the microfabrication steps for the flexible red LED probe, followed by thorough testing after assembly and packaging. The multicolor functionality is achieved by positioning the red flexible probe on top of the silicon probe, allowing the red micro-LEDs to sit above the blue LEDs. This arrangement enables the blue LEDs to emit light through the polyimide layers, either by placing the LEDs side-by-side or vertically aligned.

The design incorporates polyimide openings to accommodate the recording electrodes, ensuring accurate neuronal signal acquisition with adequate alignment margins. A significant challenge addressed in this study is the complete passivation and isolation of the red micro-LEDs and their driving metal traces, a task not previously reported. Polyimide, along with the polyimide-based wafer bonding material HD3007, is utilized for its stability throughout the multi-temperature and pressure fabrication processes.

The fully assembled DuoLite devices demonstrate reliable performance in terms of I-V-P characteristics for both LED types and impedance of the recording electrodes. The final device has been validated and is ready for in-vivo testing, as shown in experiments with a mouse model equipped with two types of viruses for blue and red light stimulation.

## Conclusion and future directions

The development of the multicolor probe required a novel process architecture to integrate red micro-LEDs into a flexible probe, using materials distinct from those in the blue-LED wafer. To achieve this, we employed polyimide and HD3007 within a microfabrication process that included both top-p-trace and bottom-n-trace fabrication steps. This approach allowed us to successfully create flexible probes with integrated red micro-LEDs for both large and small area applications. These red probes were then stacked onto conventional blue LED silicon optoelectrodes, demonstrating reliable functionality for all components after full assembly. The device’s effectiveness was validated through a head-fixed acute animal experiment.

There is an increasing demand for long-term, chronic studies involving freely moving animals, especially with bidirectional optogenetic control schemes. Considering the large number of neurons in the brain and other organs, having more active sites per module would enhance the depth of neuroscience research. Thus, advancing research on high-density multicolor neural interfaces or chronic multicolor flexible probes is essential. This progress may also involve integrating compact integrated circuits (ICs) with the probe to facilitate system miniaturization.

## References

1. Deisseroth, K. Optogenetics: 10 years of microbial opsins in neuroscience. Nat Neurosci 18, 1213–1225 (2015). 10.1038/nn.4091

2. Zhao, S., Ting, J., Atallah, H. et al. Cell type–specific channelrhodopsin-2 transgenic mice for optogenetic dissection of neural circuitry function. Nat Methods 8, 745–752 (2011). 10.1038/nmeth.1668

3. Deisseroth, K. Optogenetics. Nat Methods 8, 26–29 (2011). 10.1038/nmeth.f.324

4. Dufour, S., Koninck, Y. D. Optrodes for combined optogenetics and electrophysiology in live animals. Neurophotonics 2, 031205 (2015) 10.1117/1.NPh.2.3.031205

5. Jong, L. W., Nejad, M. M., Yoon, E., Cheng, S., Diba, K. Optogenetics reveals paradoxical network stabilizations in hippocampal CA1 and CA3. Current Biology 33, 1689-1703.e5 (2023). 10.1016/j.cub.2023.03.032

6. Wang, J., Wagner, F., Borton, D. A., Zhang, J., Ozden, I., Burwell, R. D., Nurmikko, A. V., Wagenen, R. V., Diester, I., Deisseroth, K. Integrated device for combined optical neuromodulation and electrical recording for chronic in vivo applications 9, 016001 (2011). 10.1088/1741-2560/9/1/016001

7. Wu, F. Stark, E. Ku, P.-C. Wise, K.D. Buzsáki, G. and Yoon, E. Monolithically integrated µLEDs on silicon neural probes for high-resolution optogenetic studies in behaving animals. Neuron 88, 1136–1148 (2015).

8. Buzsáki, D. Stark, E. Berényi, A. Khodagholy, D. Kipke, D.R. Yoon, E. Wise, K.D. Tools for probing local circuits: high-density silicon probes combined with optogenetics. Neuron 86, 92–105 (2015).

9. Valero, M. Zutshi, I. Yoon, E. and Buzsáki, G. Probing subthreshold dynamics of hippocampal neurons by pulsed optogenetics. Science 375, 6580 (2022).

10. Jong, L. W. Nejad, M.M. Yoon, E. Cheng, S. and Diba, K. Optogenetics reveals paradoxical network stabilizations in hippocampal CA1 and CA3. Current Biology 33, 1689–1703 (2023).

11. K. Kampasi, E. Stark, J. Seymour, K. Na, H. G. Winful, G. Buzsáki, et al., “Fiberless multicolor neural optoelectrode for in vivo circuit analysis,” Scientific reports, vol. 6, p. 30961, 2016.

12. O. Noked, A. Levi, S. Someck, O.A.- Vitos, and E. Stark, “Bidirectional optogenetic control of inhibitory neurons in freely-moving mice,” IEEE Transactions on Biomedical Engineering, vol. 68, no. 2, 2021.

13. L. Li, L. Lu, Y. Ren, G. Tang, Y. Zhao, X. Cai, Z. Shi, H. Ding, C. Liu, D. Cheng, Y. Xie, H. Wang, X. Fun, L. Yin, M. Luo, and X. Sheng, “Colocalized, bidirectional optogenetic modulations in freely behaving mice with a wireless dual-color optoelectronic probe,” Nature Communications, vol. 13, no. 839, 2022.

14. V. Gradinaru et al., “Molecular and cellular approaches for diversifying and extending optogenetics,” Cell, vol. 141, pp. 154–165, 2010

15. V. S Sohal et al., “Parvalbumin neurons and gamma rhythms enhance cortical circuit performance,” Nature, vol. 459, pp. 698–702, 2009.

16. X. Han and E. S. Boyden, “Multiple-color optical activation, silencing, and desynchronization of neural activity, with single-spike temporal resolution,” PLOS ONE, vol. 2, 2007, Paper e299.

17. B. Y. Chow et al., “High-performance genetically targetable optical neural silencing by light-driven proton pumps,” Nature, vol. 463, pp. 98–102, 2010.

18. A. S. Chuong et al., “Channelrhodopsin-2, a directly light-gated cation-selective membrane channel,” Proc. Nat. Acad. Sci., vol. 100, pp. 13940–13945, 2003.

19. G. Nagel et al., “Channelrhodopsin-2, a directly light-gated cation selective membrane channel,” Proc. Nat. Acad. Sci., vol. 100, pp. 13940–13945, 2003.

20. N. C. Klapoetke et al., “Independent optical excitation of distinct neural populations,” Nat. Methods, vol. 11, pp. 338–346, 2014.

21. Kong, D.-J. Kang, C.-M. Lee, J.-Y. Kim, J. and Lee, D.-S. Color tunable monolithic InGaN/GaN LED having a multi-junction structure. Optics Express 24, A667 (2016).

22. E. Stark, T. Koos, and G. Buzsáki, “Diode probes for spatiotemporal optical control of multiple neurons in freely moving animals,” J. Neurophysiol., vol. 108, pp. 349–363, 2012.

23. K. Fan, J. Tao, Y. Zhao, P. Li, W. Sun, L. Zhu, J. Lv, Y. Qin, Q. Wang, J. Liang, W. Wang, “Size effects of AlGaInP red vertical micro-LEDs on silicon substrate,” Results in Physics, vol. 36, no. 105449, 2022.

24. K. Kim, M. Vöröslakos, J. P. Seymour, K. D. Wise, G. Buzsáki, and E. Yoon, “Artifact-free and high-temporal-resolution in vivo opto-electrophysiology with microLED optoelectrodes,” Nature Communications, vol 11, no. 2063, 2020.

25. M. Vöröslakos, K. Kim, N. Slager, E. Ko, S. Oh, S. S. Parizi, B. Hendrix, J. P. Seymour, K. D. Wise, G. Buzsáki, A. F.-Ruiz, and E. Yoon, “HectoSTAR µLED optoelectrodes for large-scale, high-precision in-vivo opto-electrophysiology,” Advanced Science, vol. 9, no. 2105414, 2022.

26. University of North Carolina Vector Core. https://www.med.unc.edu/genetherapy/vectorcore.

27. J. E. Osborne, J. T. Dudman, PLoS One 2014, 9, e89007.

28. Pachitariu M., Steinmetz N., Kadir S., Carandini, M. & Harris, K. D. Kilosort: realtime spike-sorting for extracellular electrophysiology with hundreds of channels. bioRxiv. 10.1101/061481

29. B. R. Rost, J. Wietek, O. Yizhar, and D. Schmitz, “Optogenetics at the presynapse,” Nature Neuroscience, vol. 25, pp. 984–998, 2022.

